# New Genotypes and Diversity of *Orientia tsutsugamushi* Isolated from Scrub Typhus Patients in South Korea as Determined by Multilocus Sequence Typing

**DOI:** 10.1101/2020.08.13.249219

**Authors:** Joo-Hee Hwang, Jeongsik Kim, In O Sun, Tae Hee Lee, Kyung Min Chung, Chang-Seop Lee

## Abstract

*Orientia tsutsugamushi*, an obligate intracellular organism, is the causative agent of scrub typhus, which is endemic in the Asia-Pacific region. No comparative studies on the genotypic properties of *O. tsutsugamushi* have been performed using multilocus sequence typing (MLST) in South Korea. Here, we characterized 51 clinical isolates from Jeonju, in southwestern Korea, and we compared them to isolates from Thailand, Laos, and Japan. We also identified 10 new alleles and six novel sequence types. Overall, our results suggest that the relative genetic stability and the clonal populations of *O. tsutsugamushi* strains in South Korea have remained mostly conserved.

**Author summary:** Scrub typhus is a life-threatening disease, caused by infection with *O. tsutsugamushi*, a Gram-negative intracellular bacterium. Approximately one million people are infected globally every year, especially in the Asia-Pacific region. Strains of *O. tsutsugamushi* are typically distinguished serologically on the basis of sequences of the highly polymorphic 56-kDa outer membrane protein. Multilocus sequence typing (MLST) is a generic typing method that provides a unified bacterial isolate characterization approach that can be used for evolutionary and population studies of bacteria. In this study, we describe the development and application of a MLST scheme that was applied to 51 *O. tsutsugamushi* isolates. We found 10 new alleles and six new STs, which yielded a total of seven *O. tsutsugamushi* STs in South Korea. Among seven different STs (ST 48, 93-98), ST 48 account for the largest proportion (49.0%) of *O. tsutsugamushi* STs in South Korea. With the exception of the appearance of six novel STs, the clonal populations have remained conserved but further study of population structure and evolutionary trends is warranted.

## Introduction

Scrub typhus is an acute febrile illness caused by *Orientia tsutsugamushi*, which is transmitted by a chigger bite [1]. The clinical course of scrub typhus can range from a self-limiting to a fatal illness with an estimated mortality of 6.0% (median, range 0%–70.0%) [2]. Disease severity appears to be strain dependent, although the basis for this is unknown and is multifactorial [3,4].

*O. tsutsugamushi* strains are usually distinguished serologically on the basis of sequences of the type-specific antigen (TSA) gene that encodes the highly polymorphic 56-kDa outer membrane protein. However, use of a single surface protein gene could result in low resolution or may provide misleading evidence on evaluation of strain relatedness. Multilocus sequence typing (MLST) is a generic typing method that provides a unified bacterial isolate characterization approach that can be used for evolutionary and population studies of bacteria, regardless of their diversity, population structure, or evolution. The MLST scheme for *O. tsutsugamushi* isolated in Thailand was first proposed in 2010 [5]; in the past decade, only two studies have been conducted that used an MLST approach for *O. tsutsugamushi* typing; they evaluated isolates from Cambodia and Laos [4,6].

Here, using an MLST approach, we present the results of a prospective study to determine the geographic structure of *O. tsutsugamushi* isolated from scrub typhus patients in South Korea and we conducted a comparative analysis with previous data collected from Thailand, Laos, and Japan.

## Methods

### Patients and data collection

A prospective study was conducted in two tertiary hospitals in Jeonju, in southwestern Korea, between January 2016 and December 2017. Patients ≥18 years old who were clinically suspected of having scrub typhus were eligible. Laboratory diagnosis of scrub typhus was made according to one of the following criteria: (1) increase in indirect immunofluorescence assay (IFA) IgM titer ≥1:160 against *O. tsutsugamushi*; (2) increase in IFA IgG titer ≥1:256; (3) ≥4-fold increase in IFA titer in paired sera; and (4) positive result from nested polymerase chain reaction (PCR) targeting the 56-kDa TSA gene of *O. tsutsugamushi*.

### DNA extraction from scrub typhus patients

Whole blood samples from scrub typhus patients were collected and peripheral blood mononuclear cells (PBMC) were isolated using Lymphoprep^™^ density gradient medium and SepMate™ tubes (Stemcell Technologies, Vancouver, Canada). The isolated PBMC were aliquoted and stored at −80°C.

After isolation, DNA was purified using the QIAamp DNA Mini Kit (QIAGEN GmbH, Hilden, Germany) according to the manufacturer’s instructions [7]. Purified DNA was aliquoted and stored at −20°C.

### DNA amplification and sequencing for bacterial identification

To confirm the presence of *O. tsutsugamushi*, a nested PCR targeting the 56-kDa gene of *O. tsutsugamushi* was performed. Primers 34 (forward, 5’-TCA AGC TTA TTG CTA GTG CAA TGT CTGC-3’; the 56-kDa gene based on the Gilliam strain) and 55 (5’-AGG GAT CCC TGC TGC TGT GCT TGC TGCG-3’) were used in the first PCR. Nested PCR primers 10 (5’-GAT CAA GCT TCC TCA GCC TAC TAT AAT GCC-3’) and 11 (5’-CTA GGG ATC CCG ACA GAT GCA CTA TTA GGC-3’) were used in the second PCR amplification to generate a 483-bp fragment. Nested PCR was performed as described by Lee et al.[8]. The amplified PCR products were confirmed by agarose gel electrophoresis and purified from the agarose gel using the QIAquick gel extraction kit (QIAGEN) and the Expin™ Gel SV kit (GeneAll, Seoul, Korea). Each PCR product was analyzed and confirmed by sequencing.

### Multilocus sequence typing (MLST) and data analyses

Genomic DNA of *O. tsutsugamushi* was characterized using MLST as previously described [5]. Housekeeping genes (*gpsA, mdh, nrdB, nuoF, ppdK, sucB, sucD*) were amplified by PCR and sequenced in forward and reverse directions with their associated primers (Supplementary Table S1). The amplified PCR products were confirmed by agarose gel electrophoresis, purified, and sequenced.

### Sequence data and chromatograms were edited and analyzed using Bioedit

7.0.4. Allele numbers were assigned by querying the PubMLST database, and new alleles and profiles were submitted to the PubMLST database to obtain new allele and ST numbers (http://pubmlst.org/otsutsugamushi/). The relationships between sequence types (STs) were visualized using goeBURST [9]. The ratio of non-synonymous to synonymous nucleotide substitutions (dN/dS) of the partial sequences of seven housekeeping genes were analyzed and calculated using START2 (http://pubmlst.org/software/analysis/start2/) [10]. The phylogenetic tree of the STs was constructed based on 2,700 base pairs (bp) from concatenated sequences of all genes analyzed in specific order—*gpsA-mdh-nrdB-nuoF-ppdK-sucB-sucD*—using the neighbor-joining tree-building algorithm implemented in MEGA X [11]. We calculated the F-statistic (F_ST_) value, which indicates the degree of genetic differentiation and gene flow among populations from different geographical locations [12]. The index of diversity (D), as defined by Simpson [13], was estimated using a previously published formula [14]. The average pairwise nucleotide difference/site (Pi(π)/site), average pairwise nucleotide difference/sequence (*K*), and Tajima’s *D* value were estimated based on the new STs identified in this study using DnaSP 6.12.03 [15].

### Statistical analyses

Descriptive statistics were expressed as numbers and percentages, means and ranges, or medians and interquartile ranges (IQR). Statistical analyses were performed with MedCalc for Windows, version 19.3 (MedCalc Software, Mariakerke, Belgium).

### Ethical statement

This study was approved by the Institutional Review Board (IRB) of Jeonbuk National University Hospital, and all patients provided written informed consent (IRB registration number 2019-08-046).

## Results

### Clinical characteristics of patients with scrub typhus

Among the 51 patients, 32 (62.7%) were female, and mean age was 65.84±15.02 years. The majority of patients presented with an eschar (96.1%) or a skin rash (88.2%). A majority of patients had a headache and developed gastrointestinal symptoms. In laboratory findings, almost all patients showed mild to moderate enzyme elevation in their liver-function tests, and most patients also had thrombocytopenia. All patients were successfully treated with doxycycline or azithromycin. Patients’ demographic and clinical characteristics are summarized in Table 1.

### Bacterial isolation and diversity

A total of 51 patients’ samples that were confirmed for *O. tsutsugamushi* by nested PCR underwent MLST analysis: 51 of the 57 samples (89.5%) were successfully amplified and sequenced, while the remaining six did not yield sufficient quality data.

MLST revealed 10 new alleles in the housekeeping genes: *gpsA*, n=2; *mdh*, n=1; *nrdF*, n=1; n*uoF*, n=1; *ppdK*, n=1; *sucB*, n=2, and *sucD*, n=2. These novel alleles led to assignment of six new STs (STs 93–98; Table 2). The 51 isolates corresponded to seven different STs (ST 48, 93–98), with ST 48 accounting for the largest proportion (49.0%) of *O. tsutsugamushi* STs in Jeonju, South Korea (Table 2). ST 93 (41.2%) was the second-most-common ST and the remaining five STs (STs 94–98) were represented by a single isolate each.

The South Korea isolates had a relatively low degree of genetic diversity with a Simpson’s index of diversity of 0.58 (95% CI: 0.50–0.66) compared with the index value (0.95; 95% CI: 0.94–0.96) for all isolates including Thailand and Laos [5,6].

### Phylogenetic analysis of *O. tsutsugamushi*

To estimate the phylogeny of *O. tsutsugamushi*, we included seven STs from South Korea including ST 48/Boryong, 87 STs from Thailand and Laos, and four reference strains (Gilliam, Kato, Karp, Ikeda), for a total of 98 strains. The neighbor-joining tree showed the phylogenetic relationship among the 98 STs (Figure 1). Interestingly, the South Korea isolates were clustered in two different clades, and the five Korean STs (48, 93, 94, 95, and 98) were genetically close to ST 80 that was isolated from Laos. The remaining two STs (96 and 97) were genetically close to ST 49 (Ikeda, Japan; Figure 2).

**Figure 1.**
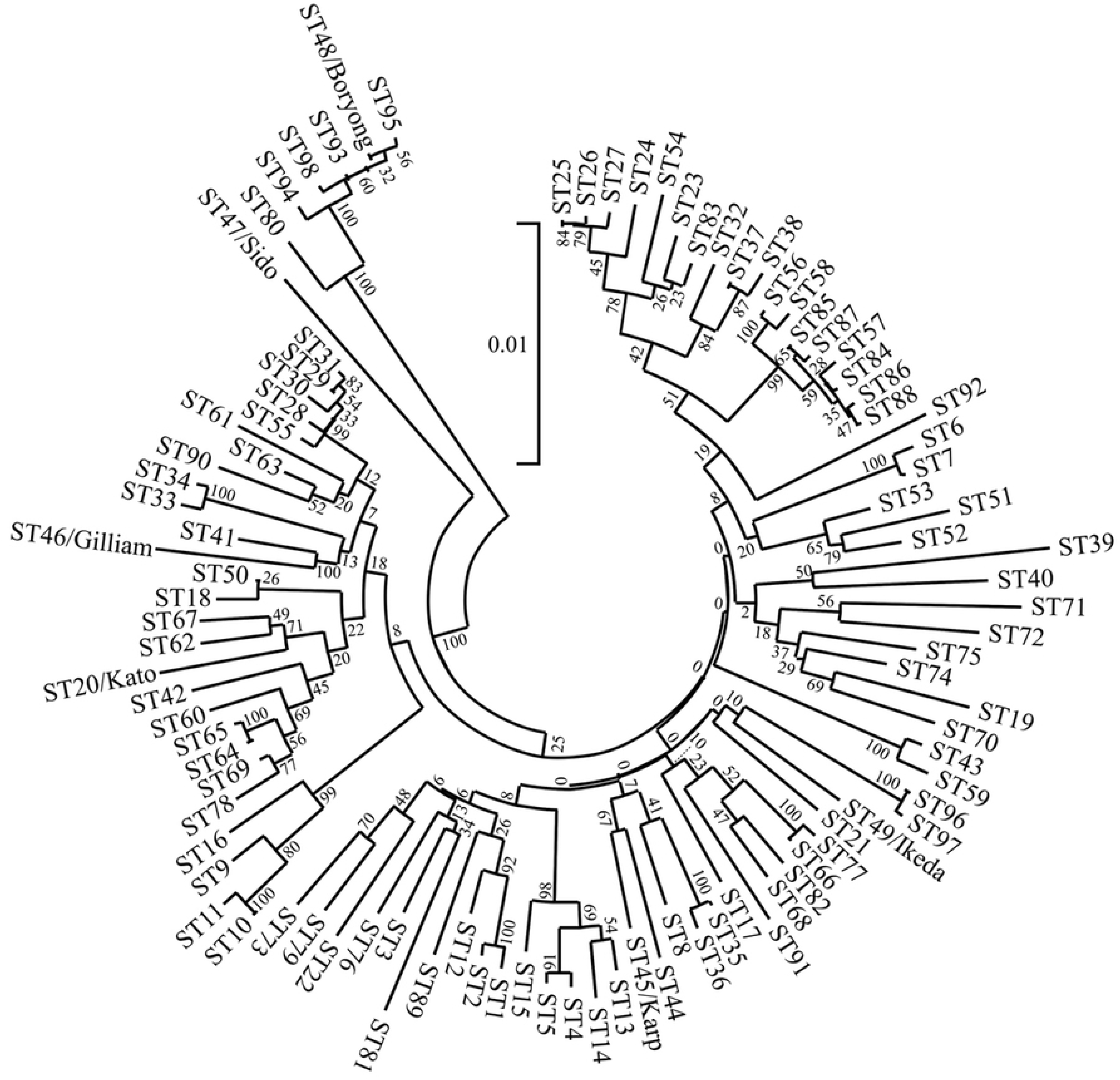
Phylogenetic analysis of 98 *O. tsutsugamushi* strains.

**Figure 2.**
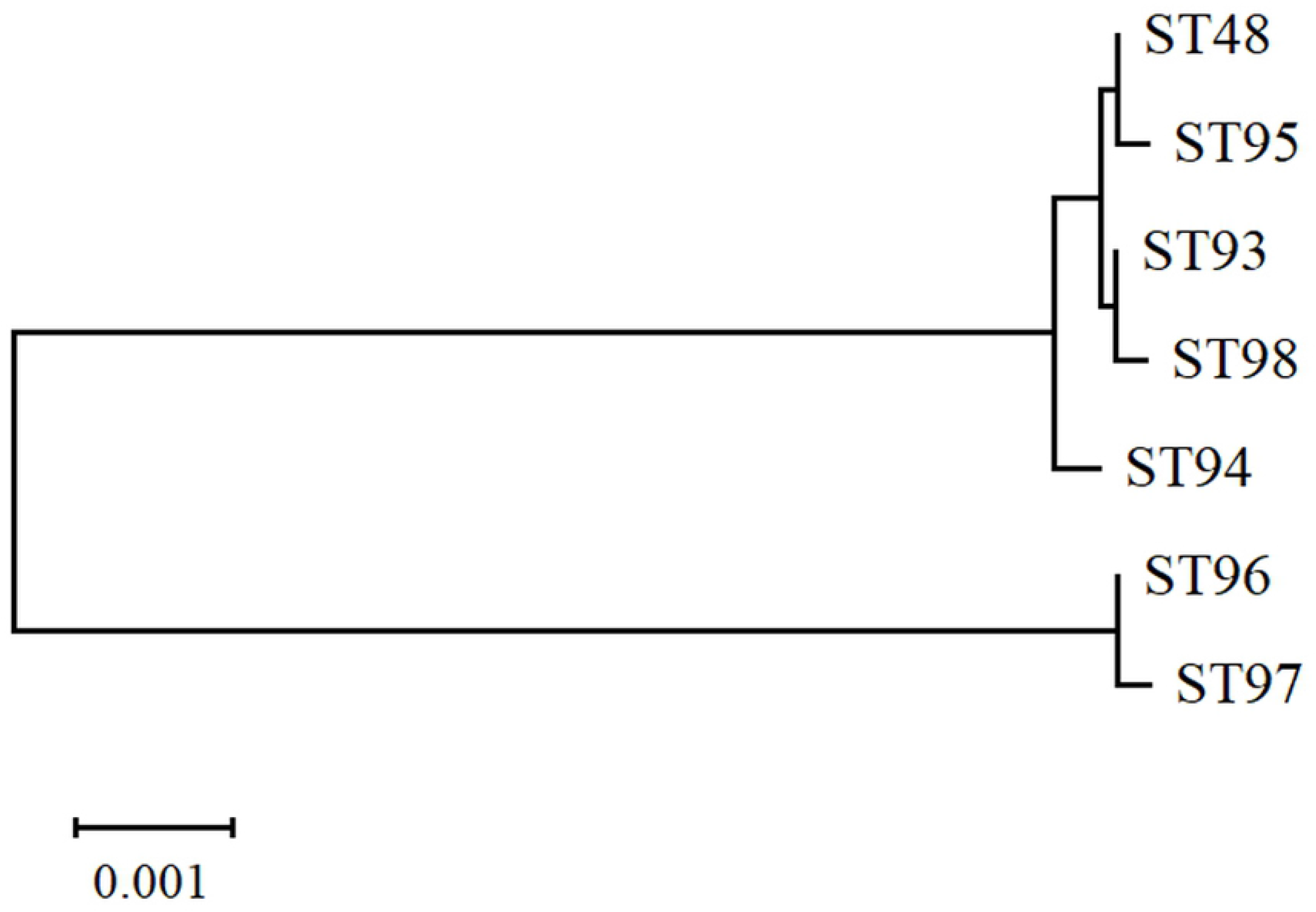
Phylogenetic analysis of 7 *O. tsutsugamushi* strains in South Korea.

Pairwise F_ST_ values for South Korea, Thailand, Laos, and Japan were calculated. The F_ST_ values ranged from 0.0215–0.4087. The neighbor-joining tree for the four populations revealed four clusters (Figure 3). These results suggest there are relatively high levels of genetic differentiation between the populations of South Korea and Thailand (0.41), Laos (0.40), and Japan (0.42).

**Figure 3.**
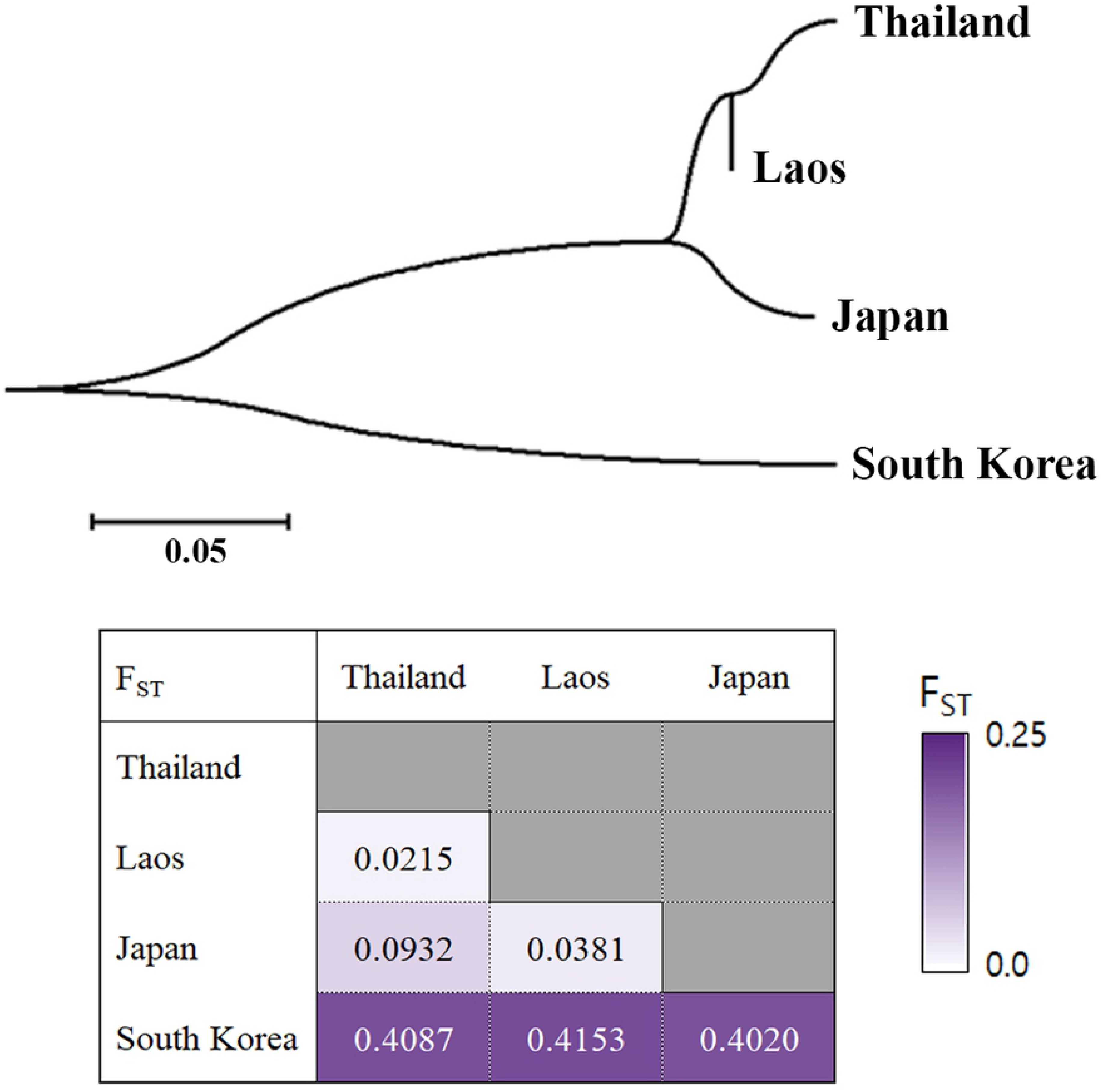
Population differentiation based on concatenated sequences of 7 housekeeping genes.

### Population structure of *O. tsutsugamushi*

Evolutionary trends and phylogenetic relationships among STs from 51 South Korean isolates and from Thailand and Lao isolates were visualized using goeBURST (Figure 4). Only one clonal complex (CC) was identified: ST 48, which is the most common, defined ST 93 and 95 as single locus variants (SLVs) using a stringent definition of 6/7 shared alleles and ST 98 as a double locus variant (DLV). There was almost no overlap among the isolates from the three countries reported above.

**Figure 4.**
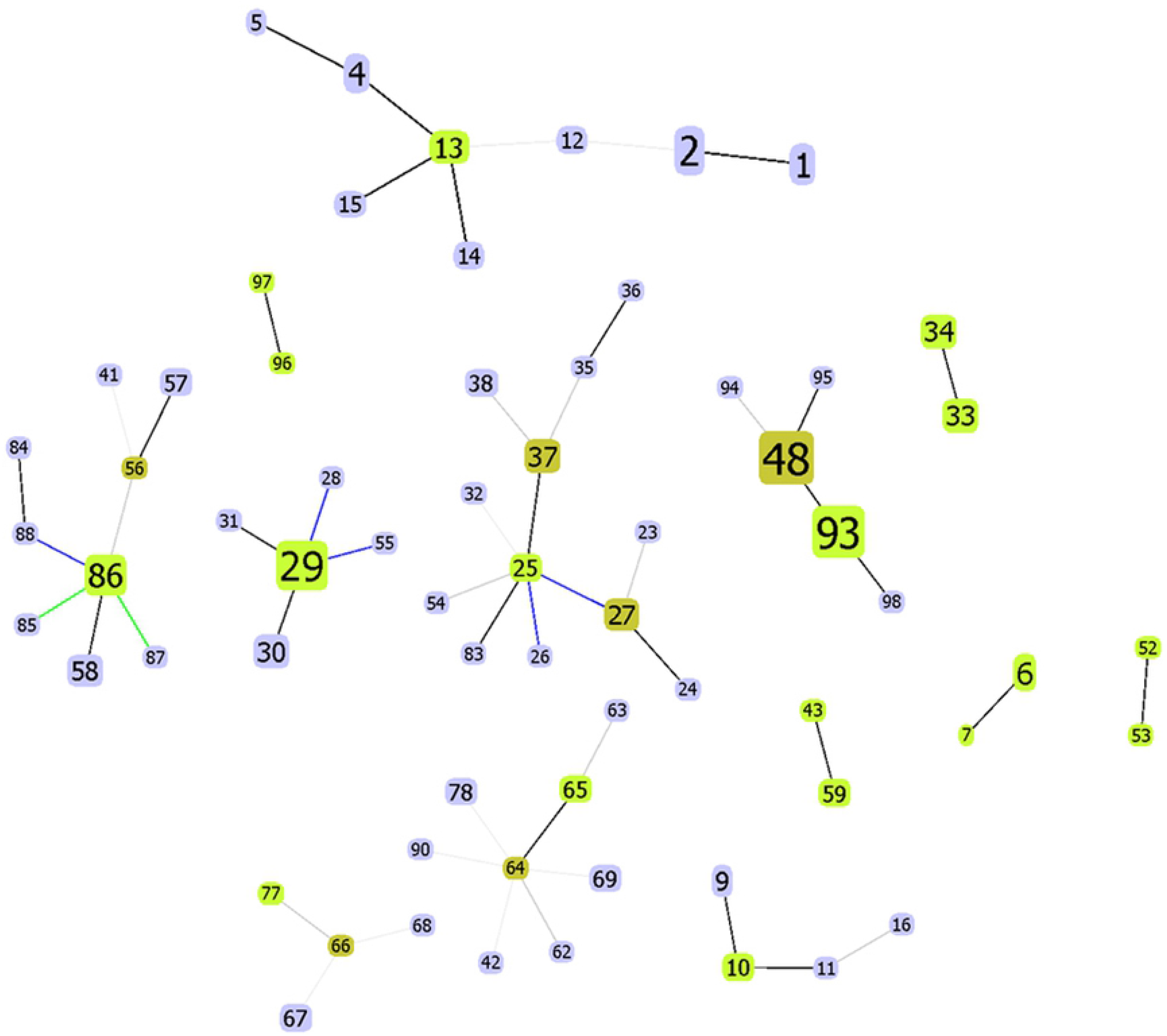
Clonal complex of the *O. tsutsugamushi* population using geoBURST.

### Descriptive analysis of nucleotide sequence data

MLST analyses of the South Korea isolates identified 10 new alleles in housekeeping genes: *gpsA* loci 43 and 44; *mdh* locus 27; *nrdB* locus 35; *nuoF* locus 42; *ppdK* locus 39; *sucB* loci 29 and 30; and *sucD* loci 37 and 38. Alleles varied from 7.5% for *mdh* to 15.4% for *gpsA* (Table 3). The MLST dendrogram generated by concatenating the seven housekeeping gene sequences (2,700 bp) showed 288 polymorphic sites. Regarding our findings, moderate nucleotide diversity of all sequenced genes was observed (π = 0.01 for *mdh*, *nrdB*, *sucB*, and *sucD*; 0.02 for *gpsA*; 0.03 for *nuoF* and *ppdK*) (Table 3).

Tajima’s *D* value is a widely used test of neutrality in population genetics. In this study, Tajima’s *D* value for the dataset containing all *O. tsutsugamushi* isolates yielded negative values, which indicate a high level of low frequency polymorphisms compared with the expected level in a neutral model.

The ratio of dN/dS was much lower than for the seven housekeeping genes, and it varied from 0.0011 for *nrdB* to 0.2790 for *sucD* (Table 3). In the South Korea isolates, the dN/dS ratio corresponded to a range of 0.0000–0.3593 (Table 4). These results suggest that all the housekeeping genes are predominantly evolving by purifying selection. These also showed that synonymous substitutions predominated over nonsynonymous substitutions for all *O. tsutsugamushi* genes we tested. We noticed that the dN/dS ratios for *nrdB* and *sucD* were lower than for the other genes. These results indicate that housekeeping genes are under strong selective constraint in the *O. tsutsugamushi* genome.

## Discussion

In this study, we explored the genetic diversity of *O. tsutsugamushi* strains isolated from scrub typhus patients in South Korea. We found 10 new alleles and six new STs, which yielded a total of seven *O. tsutsugamushi* STs in South Korea. We evaluated genetic relatedness among the South Korea *O. tsutsugamushi* strains and found that they represent a relatively distinct group composed of two subclades. This is the first report of the genetic diversity of *O. tsutsugamushi* in South Korea using MLST.

ST 48 (49.0%) and ST 93 (41.2%), the two most prevalent STs from South Korea, have not been identified elsewhere and are not closely related to isolates from Thailand and Laos (Table S2 and Figure 3) [5,6]. This is evidenced by the pairwise F_ST_ values that revealed relatively high levels of genetic differentiation between the South Korea population and the Thailand, Laos, and Japan populations. However, we cannot determine from these results whether genetic diversity was significantly correlated with geographic distance. Rather, there may have been different selective pressures on the South Korea population. Nevertheless, interregional migration of reservoir hosts could change the distribution of genetic variation within the population in the near future. Each of the MLST genes showed low dN/dS ratios (<1, particularly *nrdB* and *sucD*), which is indicative of purifying selection, and is consistent with previous studies [5,6]. No direct association was found between STs and patients’ clinical outcomes.

When we compared the 56-kDa gene sequence to the MLST data, the 56-kDa gene sequence data were generally poorly congruent to the MLST data. While our MLST results resolved the 51 isolates into seven STs, the 56-kDa gene sequence data assigned only two antigenic types (Boryong and Karp). This suggested the MLST has higher discrimination power than single-locus typing. MLST offers a better understanding of the evolutionary patterns and pathogenesis of *O. tsutsugamushi*.

Given that our isolates were collected from a single geographic region in South Korea, our results should be interpreted with caution and not be generalized. Nevertheless, these results have important clinical implications. By performing MLST for *O. tsutsugamushi*, we found one or two allelic types per locus analyzed, resulting in a total of six novel STs. Furthermore, *O. tsutsugamushi* isolated from South Korea has a unique profile which is not found in Thailand or Laos, or other reference countries.

In conclusion, this study demonstrated the relative genetic stability of *O. tsutsugamushi* strains in South Korea over the study period. With the exception of the appearance of six novel STs, the clonal populations have remained conserved but further study of population structure and evolutionary trends is warranted.

## Acknowledgments

This work was supported by the Basic Science Research Programs (NRF-2018R1D1A3B07049557) of the National Research Foundation of Korea, which are funded by the Ministry of Education and Fund of Biomedical Research Institute, Jeonbuk National University Hospital. This research was also supported by the 2019 Scientific Promotion Program funded by Jeju National University.

